# Breaking the limit of 3D fluorescence microscopy by dual-stage-processing network

**DOI:** 10.1101/435040

**Authors:** Hao Zhang, Yuxuan Zhao, Chunyu Fang, Guo Li, Meng Zhang, Yu-Hui Zhang, Peng Fei

**Affiliations:** School of Optical and Electronic Information-Wuhan National Laboratory for Optoelectronics, Huazhong University of Science and Technology, Wuhan, 430074, China; Britton Chance center for Biomedical Photonics, Wuhan National Laboratory for Optoelectronics, Huazhong University of Science and Technology, Wuhan 430074, China; MoE Key Laboratory for Biomedical Photonics, Huazhong University of Science and Technology, Wuhan, 430074, China

## Abstract

Although three-dimensional (3D) fluorescence microscopy is an essential tool for life science research, the fundamentally-limited optical throughput, as reflected in the compromise between speed and resolution, so far prevents further movement towards faster, clearer, and higher-throughput applications. We herein report a dual-stage mutual-feedback deep-learning approach that allows gradual reversion of microscopy degradation from high-resolution targets to low-resolution images. Using a single blurred-and-pixelated 3D image as input, our trained network infers a 3D output with notably higher resolution and improved contrast. The performance is better than conventional 1-stage network approaches. It pushes the throughput limit of current 3D fluorescence microscopy in three ways: notably reducing the acquisition time for accurate mapping of large organs, breaking the diffraction limit for imaging subcellular events with faster lower-toxicity measurement, and improving temporal resolution for capturing instantaneous biological processes. Combining our network approach with light-sheet fluorescence microscopy, we demonstrate the imaging of vessels and neurons in the mouse brain at single-cell resolution and with a throughput of 6 minutes for a whole brain. We also image cell organelles beyond the diffraction limit at a 2-Hz volume rate, and map neuronal activities of freely-moving *C. elegans* at single-cell resolution and 30-Hz volume rate.

## 1. INTRODUCTION

A recurring trend in biology is the attempt to extract ever more spatial information from 3D biological specimens in an ever shorter time. Many millisecond-duration dynamic cellular processes, such as functional activities occurring in live animals, require high-speed capture of transient spatial patterns at high resolution. Massive cellular details distributed across very large three-dimensional tissue volumes also need to be obtained within an acceptable time. These quests pose substantial challenges to current 3D light microscopy[1-3]. In conventional epifluorescence microscopy such as confocal[4] and two-photon excitation microscopy[2, 5], and in the newly-emerged light-sheet fluorescence microscopy (LSFM)[3, 6-11], the optical information that these imaging techniques can provide within a certain time, known as the optical throughput, remains unsatisfactory. For example, these microscopy techniques are still incapable of offering subcellular lateral resolution of mesoscale tissues in a single acquisition across a large field-of-view (FOV), nor can they break the diffraction limit for resolving organelles within a single cell. Axially scanned light-sheet microscopy (ASLM) scanned along the propagation direction can extend the imaging FOV while maintaining a thin light-sheet, thereby greatly improving the axial resolution[12, 13]; however, it still requires the recording of many planes to create a 3D image with high-axial resolution, and transient temporal profiles may be lost because of the extended acquisition time. To improve the temporal resolution for capturing highly dynamic biological processes with light field microscopy (LFM), it is possible to retrieve the transient 3D signal distribution through post-processing of a 2D light field image recorded by a single camera snapshot, but this inevitably suffers from compromised spatial resolution and the presence of reconstruction artefacts. Other approaches include various spatial resolution enhancement techniques such as Fourier ptychographic microscopy[14, 15], structured illumination microscopy (SIM)[16, 17], and single molecular localization microscopy[18-20], which have been developed over the past decade and become widely used in life science research. At the expense of increased acquisition time, these methods can computationally reconstruct a super-resolved 3D image based on a number of low-resolution measurements that contain correlations in the space, frequency, or spectrum domain. However, as these techniques either more quickly capture 3D dynamics in one snapshot, or achieve higher resolution by computing multiple image stacks, the optical throughput of these 3D microscopy methods remains fundamentally limited and has so far prevented more their more widespread application.

Unlike the abovementioned multi-frame super-resolution techniques, single-image-based super-resolution (SISR) methods can improve resolution without necessarily increasing imaging time, thereby improving throughput. However, these methods suffer from either poor enhancement effect[21-23] or restriction of the signal characteristics[24, 25]. The recent development of image enhancement and super-resolution based on neural networks has provided a paradigm shift in light microscopy[26-31] by directly deducing a higher-resolution image from a single input. However, despite its advances in strengthening the SISR capability and providing a high end-to-end reconstruction speed, the lack of super-resolution in three dimensions yet prevents its application to the capture of cellular events in 3D tissues at the desired high throughput. Furthermore, the quest for ever-better super-resolution effects intrinsically drives current network-based super-resolution methods towards newer structures and logic designs, to enable applications that nowadays require increasingly higher spatiotemporal performance.

In this study, we propose a novel imaging strategy based on a dual-stage-processing neural network, termed DSP-Net, which can efficiently recover a 3D fluorescence image stack with improved resolution and high signal-to-noise ratio (SNR). Unlike conventional one-stage network approaches, our DSP-Net contains two mutual-feedback sub-nets that manage the de-blurring and de-pixelization of the 3D image separately, fundamentally reversing the microscopy imaging process that transforms a high-resolution (HR) target into a blurred-and-pixelated image. Using this hierarchical architecture, the network allows progressive recovery from low-resolution blurred and pixelated images to middle-resolution aliased images, and finally to HR images of the targets. By this means, a properly-trained DSP-Net can restore abundant signal details that might be lost in conventional one-stage network approaches (Fig. 1b, Supplementary Fig. S3, S5, and S6), leading to a stronger resolution-enhancing capability. Furthermore, the end-to-end non-iterative inference by the network enables ultra-high processing throughput suitable for large-scale 3D reconstruction or 4D video processing. To demonstrate the abilities of the DSP-Net when combined with different imaging methods such as confocal microscopy, LSFM, and SIM, and to push the throughput limits of 3D fluorescence microscopy, we imaged neuron activities in freely-moving *C. elegans* across a large 3300 × 830 × 55-µm FOV at single-neuron resolution, extracting the spatiotemporal patterns of neuronal calcium signaling and tracking the correlated worm activity at a volume rate of 30 Hz, which is notably faster than achievable with conventional Bessel LSFM. We also demonstrate the 3D imaging of organelles in a single cultured *U2OS* cell at a 2-Hz volume rate and 100-nm spatial resolution, which cannot be met by conventional N-SIM, and show 3D anatomical mapping of neurons and vessels in a whole mouse brain at single-cell resolution with an ultrahigh throughput of 6 minutes per brain, which is over two-orders-of-magnitude higher than achievable with conventional organ 3D imaging methods.

**Fig. 1.**
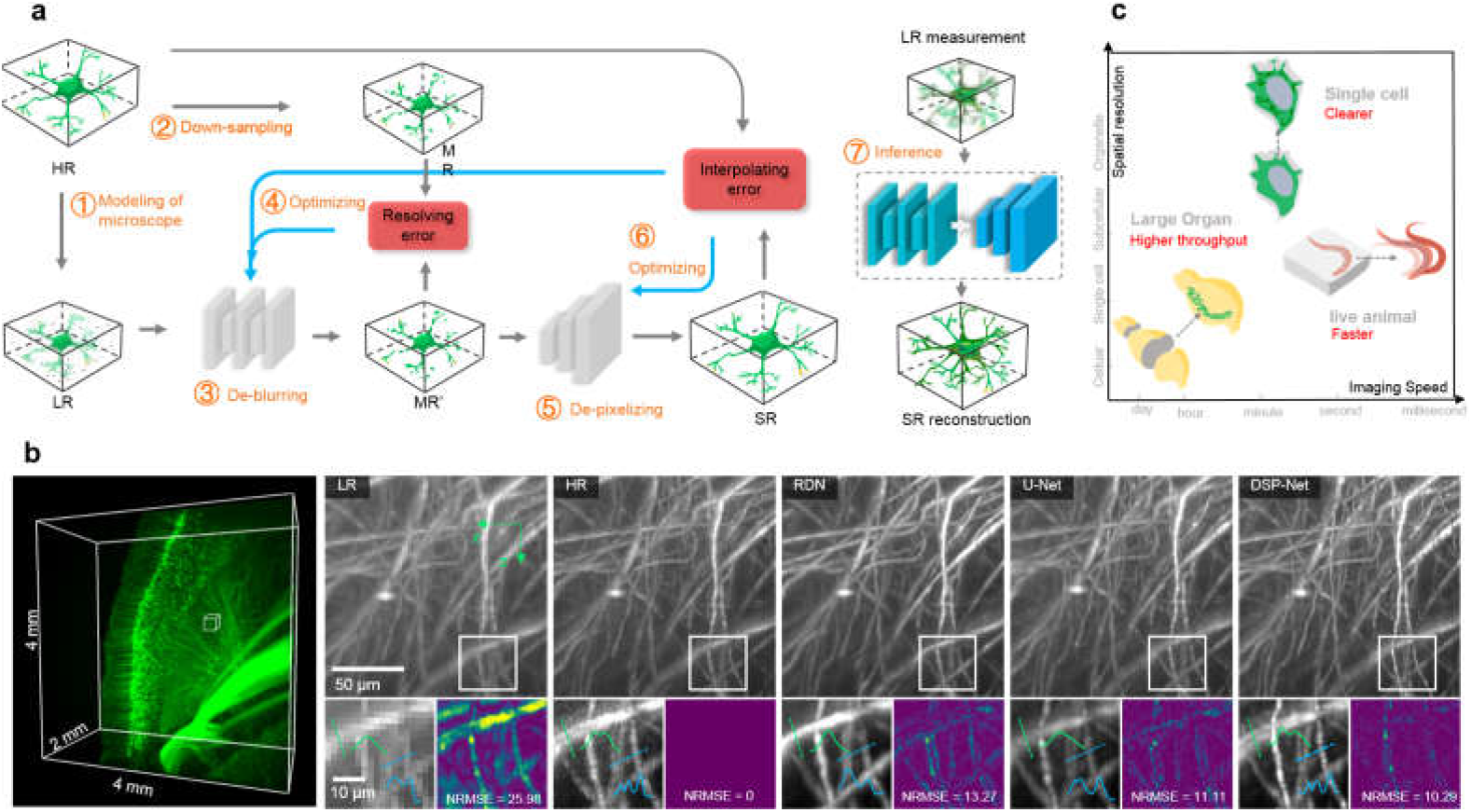
DSP-Net and its performance. **a**, The pipeline of DSP-Net. First, HR image volumes are acquired using 3D microscopy with high-magnification, high NA optics (or with 3D SRRF computation). Several number of noised, blurred, and down-sampled LR volumes are generated from these HR volumes by using a microscope degradation model (step 1). The corresponding MR volumes are simply 4-times down-sampled version of HRs (step 2), which are used as the targets in the first stage of DSP-Net, termed resolver, to restore the blurred information and reduce the noises for LR inputs (step 3). The outputs of resolver, termed MR’, are the intermediate results for computing the resolving error with the MR targets. The parameters of the resolver are iteratively optimized via minimizing the resolving error (step 4). The MR’ are further input to the second stage of the DSP-Net, termed interpolator, to recover the decimated details for MR’ (step 5). The outputs of interpolator, termed SR, are the final results for computing the interpolating error with the HR targets. At the second stage, both the interpolator (SR-HR) and the resolver (MR-MR’) are responsible for the interpolating loss, thus minimizing it would result in the optimization of both sub-nets (step 4, 6). After the DSP-Net being trained, LR images experimentally obtained by 3D microscopy with low-NA, low-magnification optics are input to the trained DSP-Net, to instantly reconstruct deblurred, denoised, and depixelated image volumes with notable quality improvement. **b**, DSP-Net enabled LSFM results of the fluorescence-labelled neurons in the mouse brain. The raw LSFM image stack (2 × 2 × 4 μm voxel) was acquired using a scanned Bessel light-sheet microscope with 3.2× magnification plus 2.7-μm thick laser-sheet illumination. The right panel shows the *xz* maximum intensity projection (1-mm projection depth) of the LR input, the HR ground truth, the SR inference by RDN, the SR inference by U-Net, and the SR inference by our DSP-Net. The line profiles of the resolved structures by each mode are plotted to indicate the achieved resolution. The normalized root mean square error (NRMSE) between each mode and the HR ground truth is calculated to evaluate the reconstruction error. **c**, The spatiotemporal map showing the DSP-Net can push the limit of conventional 3D fluorescence microscopy via enabling faster imaging speed for capturing dynamic process, higher spatial resolution for observing subcellular details beyond the diffraction limit, and higher throughput for rapidly mapping whole tissues/organs.

## 2. RESULTS

### 2.1 DSP-Net for 3D high-resolution imaging

An end-to-end deep neural network is able to accommodate complex mapping functions if it possesses sufficient parameters. However, in practice, owing to the high nonlinearity of the ill-posed SISR problem, it is often difficult to perfectly map a single low-resolution (LR) input directly to its HR target, especially when they have a significant quality gap, or in other words, when a notable quality improvement is desired. We mitigate this network limitation by logically solving the SISR problem in two steps: extracting rich details from the badly deteriorated LR input while suppressing noise and background, and enhancing the true signals. These two steps also fundamentally correspond to the restoration of signal from the optical blurring caused by an objective with a limited numerical aperture (NA) and the recovery from digital signal aliasing caused by camera pixelization. Accordingly, we created both resolver (Supplementary Fig. S1, Methods) and interpolator (Supplementary Fig. S2, Methods) modules in our DSP-Net, with each containing an independent training target corresponding to the solution of the two sub-issues. Similar to standard deep learning strategies, our DSP-Net comprises a training and an inference phase (Fig. 1a). In the training phase, the neural network establishes its resolution-enhancing ability from scratch by learning from a bunch of examples (i.e., the training dataset). HR 3D images regarded as ground truths were acquired using scanned Bessel light microscopes with high-magnification/high-NA objectives, or in the example breaking the diffraction limit, by a Bessel light-sheet microscope combined with 3D super-resolution radial fluctuations (SRRF) techniques[32]. We then applied an image degradation model (Supplementary Note S2) including optical blurring, down-sampling, and noise addition to the ground truths to generate synthetic LR images (Fig. 1a, step 1), which corresponded to our LR measurements under low-magnification/low-NA setups. Meanwhile, a directly down-scaled version of the ground truths, termed mid-resolution (MR) data, was also generated to bridge the two-stage training (Fig. 1a, step 2). With these synthetic training pairs prepared, the LR images were first fed to the resolver to clear the noise and background while retaining the potential high-frequency structures. (Fig. 1a, step 3). The mean square error (MSE) between the MR and output of the resolver, termed MR’, was defined as the resolving loss, and was used to evaluate the output quality of the resolver (Fig. 1a, step 4). The MR’ was then sent to the interpolator to be up-sampled (4 times in each dimension) into the superior-resolution (SR) results, with its high-frequency features being extracted and further enhanced (Fig. 1a, step 5). We defined the MSE between the HR ground truths and the SRs as the interpolating loss (Fig. 1a, step 6). In the first stage, the parameters of the resolver were iteratively optimized via minimization of the resolving error (step 4). In the second stage, both the interpolator and the resolver respond to the interpolating loss so that its minimization results in the optimization of both sub-nets (steps 4, and 6). By iteratively minimizing the two loss functions using a gradient descent approach, the parameters of both sub-nets can be optimized gradually, empowering the whole DSP-Net with the ability to recover SR outputs that are close to the HR ground truths corresponding to the LR inputs. Once the training of the DSP-Net converged to an optimal state, the blurred noisy pixelated 3D images captured by low-magnification/low NA (or diffraction-limited) setups could be input into the trained DSP-Net, to allow them to be directly recovered as higher-resolution, higher-contrast, de-pixelated outputs without requiring iterative computation (Fig. 1a, step 7). By substantially increasing the spatial resolution without repetitive imaging and stitching, the DSP-Net efficiently improves the imaging throughput (defined as the spatial information provided per unit acquisition time) of 3D fluorescence microscopy, which was originally limited by the system optics, regardless of the magnification factor or NA employed.

We first demonstrate the DSP-Net enabled light-sheet imaging of fluorescence-labeled neurons in the mouse brain. The raw 3D image stack, acquired using a Bessel light-sheet microscope with 3.2×/0.27 detection combined with ∼2.7-μm thickness plane illumination, encompassed 2 billion voxels across a 4 × 4 × 2-mm FOV (Fig. 1b, left). The magnified view (Fig. 1b, LR) shows obvious inadequate resolution due to the relatively low NA and large voxel size in comparison with the HR reference acquired using 12.6×/0.5 detection combined with ∼1.3-μm plane illumination (Fig. 1b, HR). The DSP-Net recovered a 3D image with an improved resolution that was very close to that of the HR reference, and also with a relatively high reconstruction fidelity (Fig. 1b, DSP-Net). Furthermore, we compared the performance of the DSP-Net with several existing one-stage pixel super-resolution / image enhancement networks[28, 33, 34]. It’s shown that the DSP-Net successfully yielded 3D images with higher resolution and fewer artefacts than one-stage networks (Fig. 1b, Supplementary Fig. S5 and S6).

### 2.2 3D imaging of a whole mouse brain at a single-neuron resolution and time-scale of minutes

We imaged GFP-tagged neurons in a whole mouse brain (Thy1-GFP-M) at high throughput. The cleared whole brain had a large size of about 10 × 8 × 5 mm, with diverse neuronal structures distributed across many brain sub-regions. Even with the use of our self-built Bessel light-sheet microscope, which has a relatively high acquisition speed (Supplementary Table S2), single-cell-resolution volumetric mapping of the whole brain remains highly challenging because of the limited system throughput, and thereby requires long-duration tile imaging, e.g., the acquisition of around 100 tiles in tens of hours under a high-magnification setup of 12.6× detection combined with 1.3-μm thin plane illumination (Methods). Furthermore, the degradation of image quality resulting from the bleaching of fluorescence and deep tissue scattering is inevitable (see both the 1st and 3rd columns of Figs. 2b2, b3, and b4).

**Fig. 2.**
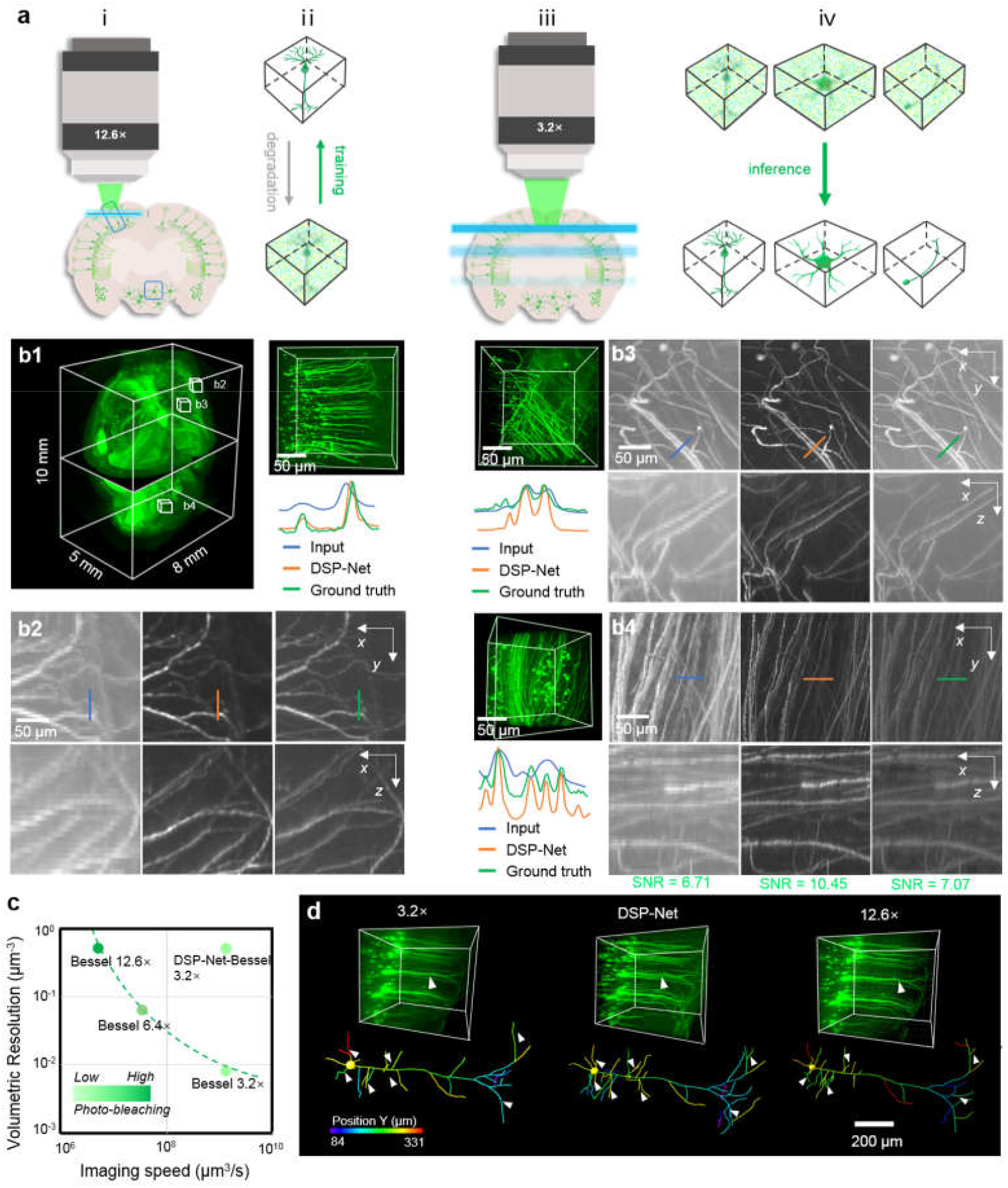
DSP-Net for 3D single-cell-resolution imaging of whole mouse brain at minutes time-scale throughput. **a**, Workflow of whole-brain data acquisition and DSP-Net processing, which includes (i) HR imaging of neurons in a few regions of the brain by 12.6× Bessel sheet (0.5 × 0.5 × 1 μm voxel, 0.1 ms exposure, 5 fps). (ii) Network training based on these HR label data. (iii) rapid imaging of neurons in the whole brain by 3.2× Bessel sheet (2 × 2 × 4 μm voxel, 0.4 ms exposure, 20 fps), with 6 tiles acquired in ∼6 minutes. (iv) DSP-Net inference to recover a digital whole brain with improved resolution (0.5 × 0.5 × 1 μm voxel) and SNR. **b**, Comparison of three ∼200 × 200 × 200 μm^3^ region-of-interests (ROIs) in isocortex (**b2**), striatum (**b3**), and cerebellum (**b4**), selected from the 3.2× DSP-Net reconstructed whole brain (**b1**, ∼10 × 8 × 5 mm^3^). The magnified views of transverse (*xy*) and coronal (*xz*) planes are shown to compare the detailed neuronal fibers resolved by raw 3.2× Bessel sheet (left), 3.2× DSP-Net (middle), and 12.6× HR Bessel sheet (right). The line profiles of the same neuron fibers indicate the axial and lateral FWHMs by 3.2× DSP-Net (orange), which are narrower than 3.2× LR inputs (blue), and close to 12.6× HR ground truths (green). Besides the resolution improvement, the low SNR caused by 20 fps acquisition rate at 3.2× (6.71 in **b4**, left) are also notably increased to a level (10.45 in **b4**, middle) higher than the SNR of 12.6× HR image with 5 fps acquisition rate (7.07 in **b4**, right). **c**, The throughput plot comparing the acquisition speed, and volumetric resolution of 3.2× (LR), 6.4×, 12.6× (HR) and DSP-Net enabled 3.2× (SR) Bessel sheet modes. The dash line connecting the data points of Bessel sheets represents the throughput limit in conventional light-sheet microscopes. **d**, Improvement for single-neuron tracing brought by DSP-Net. An identical pyramidal neuron in the isocortex region resolved by 3.2×, 3.2× DSP-Net, and 12.6× was segmented and traced using Imaris.

In our DSP-Net Bessel light-sheet implementation (Fig. 2a), we first chose high-resolution high-SNR Bessel images of a few neurons (12.6× setup, 0.4 ms exposure, 5 fps) in the isocortex and cerebellum regions to construct a training dataset for the DSP-Net, and then used the trained network to restore low-resolution low-SNR images of various types of neurons across the whole brain (3.2× setup, 0.1 ms exposure, 20 fps). In Fig. 2b, we show that the DSP-Net recovered 3D neurons from isocortical (b2, middle column), striatum (b3, middle column), and cerebellum (b4, middle column) regions of the reconstructed whole brain (b1), and compare the results with raw 3.2× inputs (left columns) and HR reference images using the same 12.6× setup used to acquire the training data (right columns). We notice that in comparison with the ambiguous LR inputs of the 3.2× Bessel sheet, the images recovered by DSP-Net show not only higher SNR, but also sharper details of the neuron fibers, providing near-isotropic 3D resolution as high as that of the reference HR 12.6× Bessel sheet. The line-cuts through individual neuronal fibers (Fig. 2b) using each method further confirm the narrower axial and lateral full width at half maximum for the 3.2× DSP-Net mode in comparison with the raw 3.2× Bessel sheet (plots in Fig. 2b). Therefore, the DSP-Net enabled Bessel light-sheet microscopy to perform rapid whole-brain imaging, typically acquiring the data within a few minutes under 3.2× acquisition at a single-neuron resolution close to that achieved by the 12.6× HR setup (Supplementary Video S1). The high throughput shown by the rapid achievement of high-resolution images across a large FOV could be over two-orders-of-magnitude higher than that achievable using conventional Bessel light-sheet microscopy (Fig. 2c, Methods). Furthermore, we demonstrate that the extra robustness of the neural network allows the recovery of various types of neurons in the whole brain while only requiring a small amount of HR neuron data for the training process. The high-throughput high-resolution mapping of the whole brain means that the DSP-Net can obviously visualize more neuron fibers (dendrites) in a pyramidal neuron than can possibly be imaged on a raw 3.2× image, thereby enabling image-based segmentation and tracing of neurons to be performed at the system level and in three dimensions (Fig. 2d).

### 2.3 Rapid 3D super-resolution imaging of a single cell with low phototoxicity

A key concern in super-resolution fluorescence microscopy is the photon consumption required for reconstructing SR images. We demonstrate the ability of the DSP-Net by imaging Alexa Fluor 488-labelled microtubules, which are beyond the diffraction limit, in single *U2OS* cells. We do this with an improved acquisition rate as well as a reduced photon budget. We first obtained an SR 3D image of microtubules based on the 3D-SRRF[32] computation of thirty diffraction-limited image stacks consecutively acquired on our custom high-magnification Bessel light-sheet microscope (Methods). Because of the noticeable photo-bleaching caused by repetitive fluorescence emission, the last (30^th^) image stack of the time series obviously shows the lowest SNR and diffraction-limited resolution, and is similar to an image stack acquired using lower-intensity laser illumination. We then used the reconstructed 3D-SRRF image (HR), its corresponding down-sampled version (MR), and a diffraction-limited measurement (LR) to construct a training dataset for the DSP-Net, and then applied the trained network to the SR reconstruction of microtubules in another *U2OS* cell (Methods, Fig. 3a). We demonstrate that for low-resolution low-SNR input data of microtubules obtained by diffraction-limited optics and low-dose excitation (emission from a low number of photons), the DSP-Net enables a notable improvement in both spatial resolution and SNR (Fig. 3c, Supplementary Fig. S8, Supplementary Video S2). We also demonstrate that the network is highly robust, being capable of recovering a high-fidelity super-resolution image stack from either the 1st or 30^th^ stack of the time-series (Fig. 3c, Supplementary Fig. S6). The line profiles of the microtubules resolved by the low-SNR inputs, high-SNR inputs, their corresponding DSP-Net outputs, and the SRRF result are plotted in Fig. 3d, allowing comparison of the resolution achieved by each method. The quality of the DSP-Net recovered results is similar to that of the SRRF results (Fig. 3d). We then repetitively imaged a cell 200 times using low-intensity plane illumination (0.3 mW at the back pupil plane) and compared its photo-bleaching rate with that occurring with the regular-intensity plane illumination used for SRRF imaging (2.1 mW at the back pupil plane). The low-intensity illumination demonstrated an obviously lower bleaching rate, which is of substantial benefit when imaging a living cell (Fig. 3e). At the same time, our DSP-Net still provided high-quality SR images based on single low-resolution low-SNR measurements (t=1, 200, respectively; Fig. 3f), while only needing around 0.5% of the photons of the SRRF imaging (Fig. 3g). We thereby demonstrate that our DSP-Net combined with rapid Bessel-sheet microscopy permits the sustained 3D imaging of cells beyond the diffraction limit at a high volume rate of about two volumes s^-1^ (camera rate 200 fps, Fig. 3h).

**Fig. 3.**
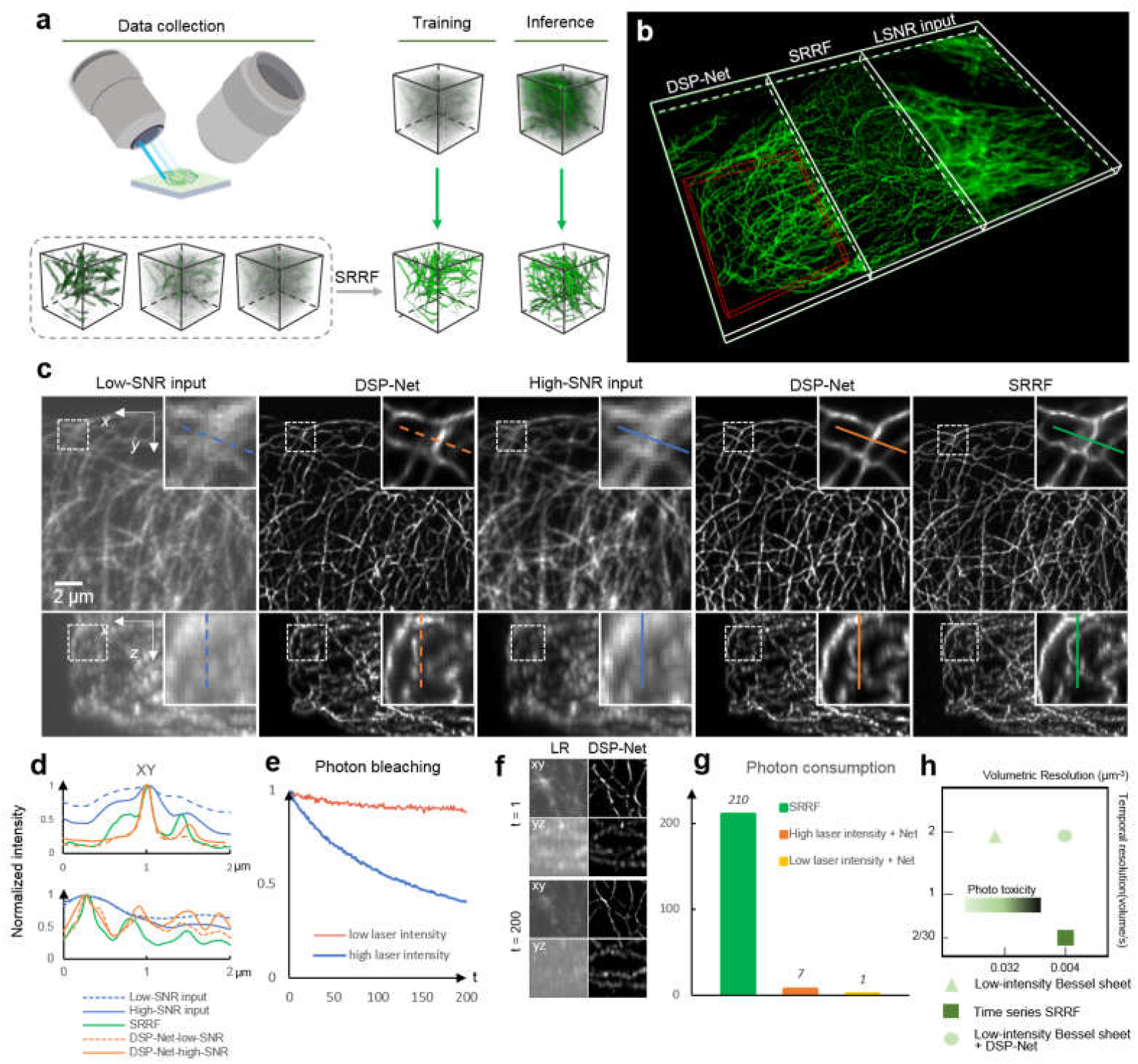
DSP-Net for 3D super-resolution imaging of antibody-labeled microtubules (Alexa-488) in single cell with high volume rate and low photon budget/toxicity. **a**, Work flow of DSP-Net 3D super-resolution imaging including: (i) SRRF reconstruction of microtubules in single cell (Alexa-488) based on 30 diffraction-limited LSFM image stacks consecutively acquired by a high-magnification Bessel light-sheet microscope (60×/1.1 detection objective plus 700 nm thickness Bessel sheet). (ii) DSP network training based on SRRF reconstructed images (HR), down-sampled SRRF images (MR), and the 30^th^ measurement with lowest SNR due to photobleaching (LR). (iii) Rapid 3D imaging of microtubules (108 × 108 × 300 nm voxel), at 2 Hz volume rate. The obtained diffraction-limited, low-SNR image stack is super-resolved by trained DSP-Net instantly. **b**, The reconstructed volume rendering of microtubules in a fixed U2OS cell, comparing the overall visual effects by diffraction-limited Bessel sheet, DSP-Net outputs and SRRF. **c**, Magnified views in *xy* and *yz* planes of the red box region in **b**. Shown from the left to right are: LR (diffraction-limited), low-SNR microtubules (30^th^ measurement); DSP-Net reconstruction of LR, Low-SNR data; LR, high-SNR microtubules (1^st^ measurement); DSP-Net reconstruction of LR, high-SNR data; and SRRF reconstruction. **d**, Line profiles of resolved microtubules by each methods, indicating their achieved lateral and axial resolutions, which are ∼110 and 270 nm for DSP-Net, respectively (FWHMs). **e**, Comparison of photo-bleaching rate of microtubules over 200 volumes acquired with different excitation intensity of 0.3 and 2.1 mW. The x-axis is the imaging times. **f**, LR measurement at t = 1 and t = 200 and the corresponding DSP-Net results. **g**, Comparison of the photon consumptions of different method for reconstructing super-resolution images. **h**, The spatial-temporal map showing the unwilling trade-off between temporal resolution, spatial resolution, and phototoxicity (triangle and square data points). DSP-Net breaks this spatial-temporal limit (circular data point).

### 2.4 Instantaneous volumetric imaging of neural activities in freely moving C. elegans

We demonstrate that DSP-Net-enabled 3D microscopy is capable of capturing dynamic processes in live model organisms by performing the imaging of neuronal activity in freely moving *C. elegans* (Fig. 4). A microfluidic chip was used to permit *C. elegans* (L4-stage) to move freely inside a large micro-chamber (3300 × 830 × 55 µm, Fig. 4b). A 4×/0.28 objective combined with a fast-scanning Gaussian light sheet (Fig. 4a, Supplementary Fig. S14) was used to rapidly image the activities of motor neurons labeled with GCaMP (strain ZM9128 hpIs595[Pacr-2(s)::GCaMP6(f)::wCherry]) at a 30-Hz volume-acquisition rate across a 3300 × 830 × 55 µm field-of-view, yielding a total of 900 volumes over a 30-second observation (Methods). Because of the use of a large-FOV objective together with a sparse scanning step (4 μm) to achieve a high recording volume rate, the raw images inevitably suffered from ambiguous cellular resolution (Fig. 4c, left), with signals from adjacent neurons remaining indiscernible. The DSP-Net then provided enhanced resolution that allowed the visualization of neuron calcium signaling during fast body movement (Fig. 4c, right, Supplementary Fig. S10). The voxel rate of the conventional 4× light-sheet was thus expanded ∼60 times by the DSP-Net (Fig. 4d), thereby achieving single-neuron resolution across the mesoscale FOV at a 30-Hz volume rate. Furthermore, the non-iterative inference of the network could sequentially enhance the 3D images at a high rate, making it suitable for processing sustained biological dynamics, which is computationally challenging using conventional methods. Based on the four-dimensional imaging results with a sufficiently high spatiotemporal resolution, we identified single specific A- and B- motor neurons that were associated with motor-program selection, and mapped their calcium activities over time (Fig. 4e, Supplementary Video S3). Meanwhile, by applying automatic segmentation of the worm body contours based on the location and morphology of neuron signals[35], we calculated the worm’s curvatures related to its locomotion and behavior, thereby allowing classification of the worm’s motion into forward, backward, or irregular crawling (Fig. 4g). The traces of transient *Ca*^2+^ signaling were found to be relevant to switches from forward to backward crawling of the worm, consistent with previous findings[36, 37] (Fig. 4e, f).

**Fig. 4.**
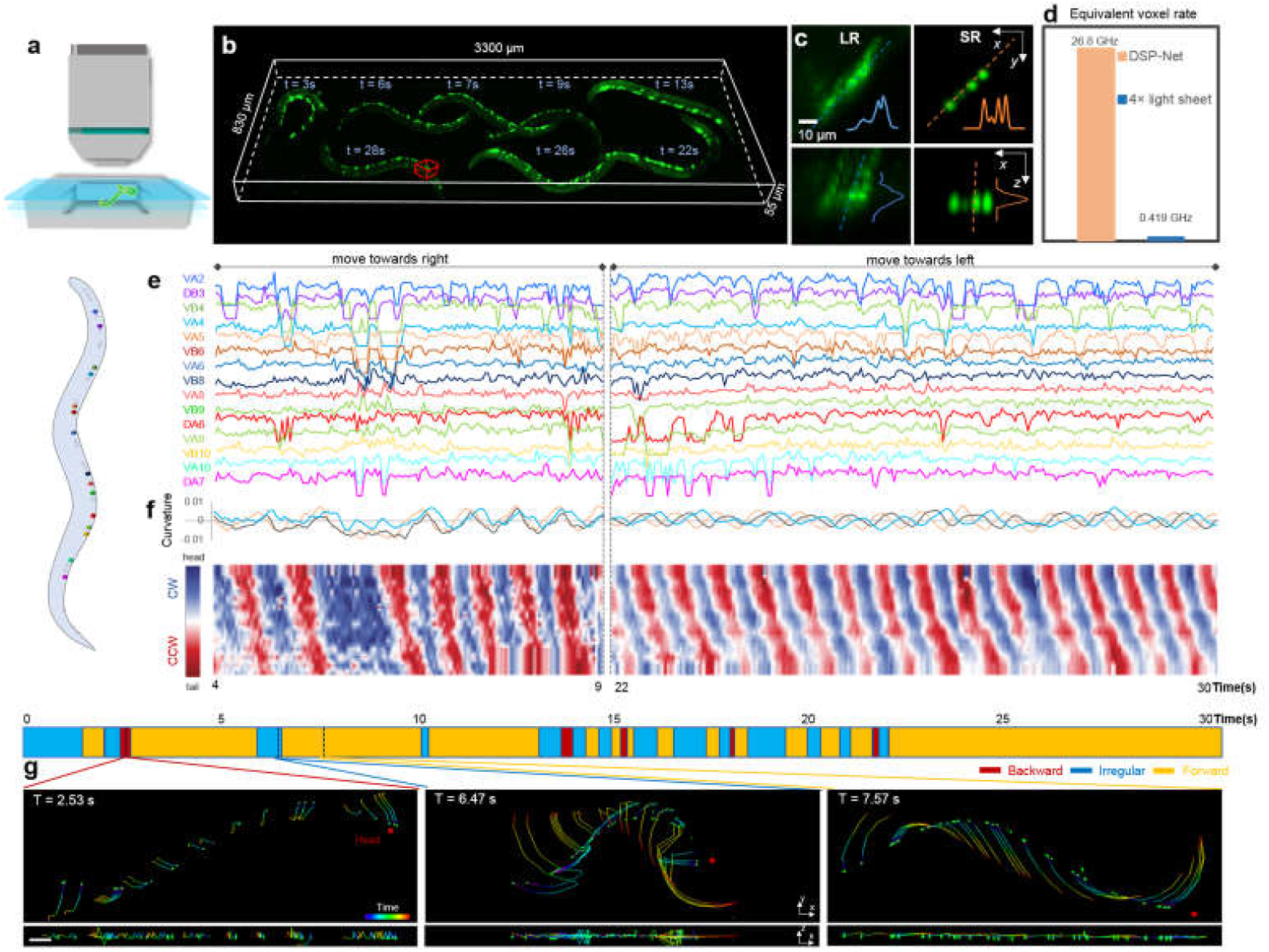
Whole-animal *Ca*^2+^ imaging of freely moving *C. elegans* using DSP-Net. **a**, Geometry combining fast light-sheet microscopy with microfluidic technique for 3D imaging of motor neuron activities (*hpls595*) of live *C. elegans* freely moving in a micro-chamber (3300 × 830× 55 µm). The acquisition rate is 30 volumes/second. **b**, The 3D trajectory of the *C. elegans* movement during a 30-seconds observation. **c**, Four neurons at one instantaneous volume (red box in **b**) by raw 4×-LSFM and after DSP-Net reconstruction, respectively. It’ obvious that DSP-Net restores single-cell resolution and reduces motion artefacts, making possible the quantitative analysis of activities from single neurons. Scale bars, 10 µm. **d**, The equivalent imaging throughput of conventional 4× light-sheet and DSP-Net-enabled 4× light-sheet. **e**, Activity of 15 identified motor neurons from the acting worm (left) by tracing the *Ca*^*2+*^ transients of individual neuron after being super-resolved by DSP-Net. Each trace indicates the fluorescence intensity change of a single neuron (Δ*F*/*F*_*0*_) during two time series of worm moving towards the right of FOV and moving back towards the left. **f**, Kymograms showing the curvature change of 14 sample points along the body of acting worm. Among them, traces of curvatures of 3 sample points are plotted on the top. The body contours of the worm are automatically obtained based on a U-net segmentation. **g**, An ethogram (top panel) based on analysis of curvature change over the entire imaging period classifies the worm behavior (lower panel) into backward (left), irregular (middle) and forward (right) statuses. Three selected volumes with time-coded traces in accordance to the ethogram visualize the forward (left, 300 ms), irregular (middle, 560 ms), and backward (right, 165 ms) crawling tendency of the worm. Scale bars, 50 μm.

### 2.5 Cross-mode 3D restoration and super-resolution by DSP-Net

We directly used a DSP-Net trained on Bessel light-sheet images of brain neurons to restore low-resolution confocal microscope images of neurons, and a DSP-Net trained on SRRF Bessel light-sheet images of microtubules in cells to restore diffraction-limited wide-field SIM microscope images of microtubules. As shown in Fig. 5a, 3D image stacks of a 100-μm thick mouse brain slice were captured by a confocal microscope (Nikon Ni-E, CFI LWD 16×/0.8 W objective). The degraded version of this confocal image stack, considered as 3.2× LR input (Fig. 5b, LR), was then input into the abovementioned DSP-Net trained on Bessel light-sheet data for cross-mode (CM) restoration. The results of this model, which we termed the CM-DSP-Net in Fig. 5b, were further compared with the recovered results of another DSP-Net that was regularly trained with confocal data (DSP-Net), and with the 12.6× (down-sampled from 16×) confocal stack (Fig. 5b, HR). For the super-resolution 3D imaging of microtubules in a *U2OS* cell (Fig. 5e), the LR input was a diffraction-limited 3D wide-field image obtained by simply averaging the fifteen SIM exposures at each *z*-plane (Nikon N-SIM, CFI Apo TIRF 100×/1.49 oil, Fig. 5f, input). The above-mentioned DSP-Net trained using Bessel light-sheet data was then applied to this wide-field image to recover 3D microtubules beyond the diffraction limit (Fig. 5f, CM-DSP-Net). The result was finally compared with the super-resolution image obtained by the 3D-SIM reconstruction using the 15 patterned exposures (Fig. 5f, 3D-SIM). In both cases, the cross-mode DSP-Net provided extraordinary enhancement of the heterogeneous inputs. Qualitative analysis of the reconstruction quality reveals the CM-DSP-Net results to show spatial resolutions similar to those of the regular DSP-Net. At the same time, the high-resolution-scaled Pearson coefficient (RSP) and low-resolution-scaled error (RSE) also indicate the sufficient accuracy of the CM-DSP-Net recovery.

**Fig. 5.**
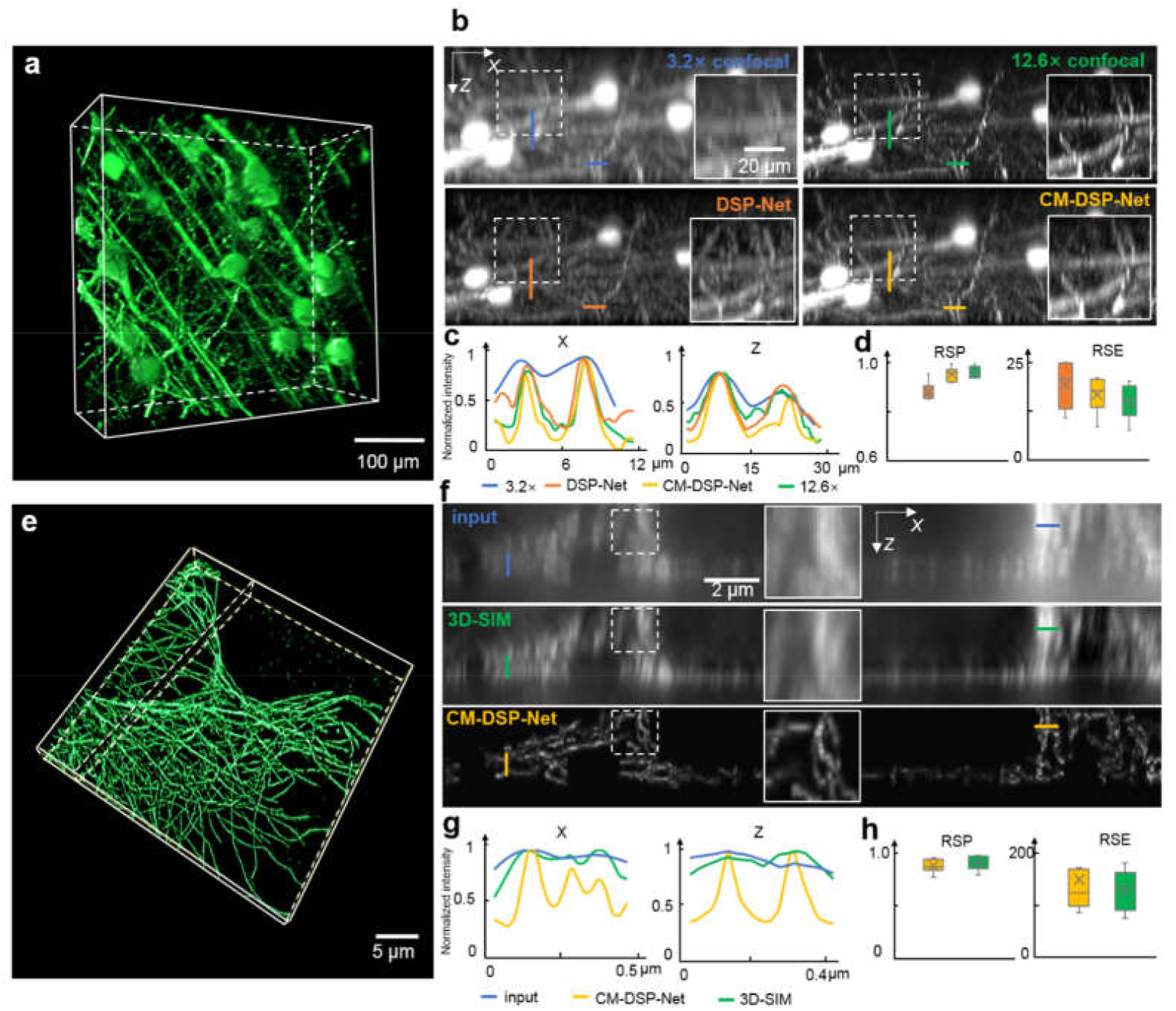
Cross-mode 3D restoration and super-resolution by DSP-Net. **a**, 3D confocal image stack of neurons (GFP) in 100-μm thick mouse brain slice recovered by cross-mode DSP-Net (CM-DSP-Net), which is trained with image data obtained by light-sheet microscope. **b**, *xz* projections of the synthetic 3.2× confocal image (blue), 12.6× confocal image (green), super-resolution of 3.2× confocal image by standard DSP-Net (yellow), and CM-DSP-Net (red), respectively. **c**, Normalized intensity plots of the linecuts through neuronal fibers along lateral (*x*) and axial (*z*) directions in **b. d**, Comparison of RSP and RSE values of the DSP-Net, CM-DSP-Net and 12.6× HR results calculated with respect to the 3.2× LR images. **e**, CM-DSP-Net reconstruction of the diffraction-limited 3D wide-field images of antibody-labelled microtubules in *U2OS* cell (Alexa-488, 100×/1.49 NA objective). The network was trained with super-resolution images by 3D-SRRF enabled Bessel light-sheet microscopy. **f**, *xz* projections of the raw measurement (blue), 3D-SIM reconstruction (green) and the CM-DSP-Net reconstruction (red), respectively. **g**, Normalized intensity plots of the linecuts through microtubules along lateral (*x*) and axial (*z*) directions in **f. h**, Comparison of RSP and RSE values of the CM-DSP-Net and 3D SIM results calculated with respect to the diffraction-limited microtubule images.

### 2.6 Cross-sample 3D restoration and super-resolution by DSP-Net

In addition to its cross-mode capability, we further demonstrate that the DSP-Net can also perform cross-sample (CS) restoration. We did this using a network trained on brain neurons at single-cell resolution to restore brain vessels, and a microtubule-trained network to perform super-resolution imaging of endoplasmic reticulum beyond the diffraction limit. In both cases, our network showed remarkable improvements for new types of signals (Fig. 6a, b, e, f, Supplementary Fig. S11). Furthermore, quantitative analyses of the achieved resolution and reconstruction errors both verify the reliability of the cross-sample recovery by DSP-Net. We note that the robust cross-mode and cross-sample capabilities shown by our 3D DSP-Net are also consistent with previous findings on 2D network super-resolution[27]. It should be noted that the CM/CS-DSP-Net can achieve resolution improvement even better than of the regular DSP-Net if the CM/CS-DSP-Net receives higher-quality training. However, the CM/CS-DSP-Net fails to generate reasonable outputs when the new data are completely different with the original data (Supplementary Fig. S11, DSP-ER recovered neurons and nuclei). In such cases, further tuning techniques such as transfer learning should be engaged.

**Fig. 6.**
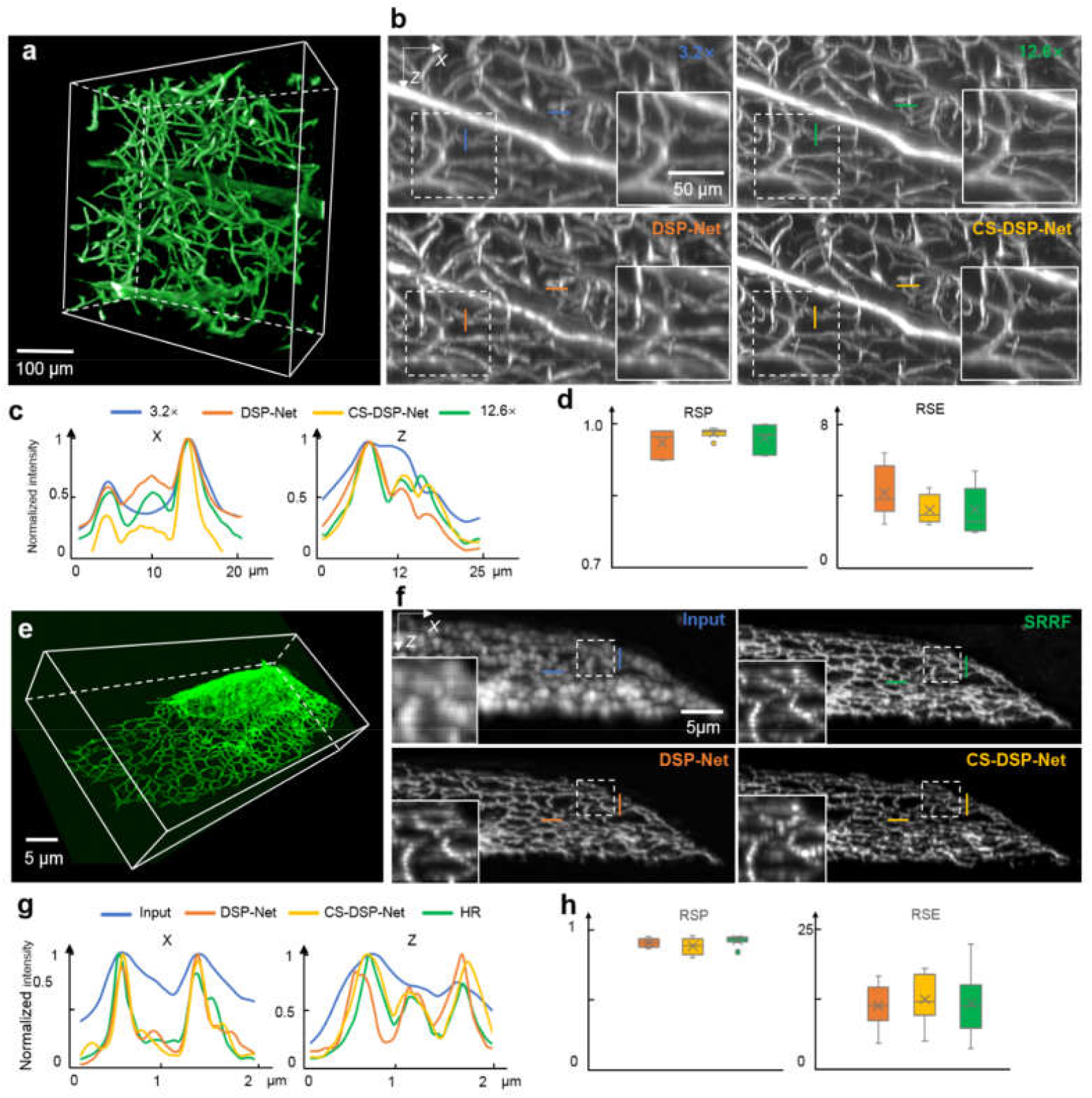
Cross-sample 3D restoration and super-resolution by DSP-Net. **a**, 3D Bessel sheet image stack of blood vessels (Alexa-647) in mouse brain recovered by cross-sample DSP-Net (CS-DSP-Net), which is trained with 3D neuron images by the same Bessel light-sheet microscope. **b**, *xz* projections of the 3.2× Bessel sheet image (blue),12.6× Bessel sheet image (green), restoration of 3.2× image by standard DSP-Net (yellow), and CS-DSP-Net (red), respectively. **c**, Normalized intensity plots of the linecuts through vessels along lateral (*x*) and axial (*z*) directions in **b. d**, Comparison of RSP and RSE values of the DSP-Net, CS-DSP-Net and 12.6× HR results calculated with respect to the 3.2× LR images. **e**, CS-DSP-Net super-resolution reconstruction of the diffraction-limited 3D wide-field images of antibody-labelled endoplasmic reticulum (the antibody labeled EGFP) in *U2OS* cell (100×/1.49 NA objective). The network was trained by microtubule images. **f**, *xz* projections of the raw diffraction-limited image of endoplasmic reticulum (blue), SRRF reconstruction (green), DSP-Net reconstruction (yellow) and CS-DSP-Net reconstruction (red), respectively. **g**, Normalized intensity plots of the linecuts through endoplasmic reticulum along lateral (*x*) and axial (*z*) directions in **f. h**, Comparison of RSP and RSE values of the DSP-Net, CS-DSP-Net and 3D-SRRF results calculated with respect to the diffraction-limited endoplasmic reticulum images.

## 3. Methods

### 3.1 Light-sheet microscopy setups

#### High-magnification Bessel-sheet microscope

Our high-magnification Bessel-sheet microscope for single-cell imaging was built based on an upright microscope (BX51, Olympus). Fiber-coupled lasers with four excitation wavelengths (405, 488, 589, 637 nm) were first collimated by a collimator (Thorlabs, F280FC-A) and then sent to an acousto-optic tunable filter (AOTF) to control their intensities. The 1^st^-order diffraction beam after AOTF was selected and expanded (Thorlabs, CBE05-A) to 10 mm diameter and then sent to an axicon (Thorlabs, AX252-A) to generate annular Bessel beam at the front focal plane of the achromatic lens (Thorlabs, AC254-075-A-ML). The interference pattern was focused onto a galvanometer (Thorlabs, GVS112), which was combined with a scan lens (Thorlabs, CLS-SL) to scan the Bessel beam into a sheet. Finally, the Bessel sheet was compressed for 20 folds through a 4-f system containing a tube lens (Thorlabs, TTL200) and an objective lens (Thorlabs, MY20×-803) to obtain the final illuminating Bessel-sheet with FWHM thickness of ∼700 nm. In the detection path of upright microscope, a water dipping objective (Olympus, LUMFLN 60XW 60×, 1.1 NA) was used to collect the fluorescence emission from the fluorophore-tagged U2OS cell. To eliminate the emission excited by the side lobes of the Bessel beam, the rolling shutter of the detection sCMOS camera (Hamamatsu, ORCA Flash 4.0 V3) was synchronized with the beam scanning to form an electronic confocal slit so that only the fluorescence excited by the central maximum of the Bessel beam could be detected. Finally, we obtained a high-quality Bessel sheet illumination with ∼700 nm uniform thickness covering the entire illuminated range of the U2OS cell. For 3D imaging, the cells attached to the cover glass was mounted onto a customized holder and vertically (*z*) scanned by a piezoelectric stage (P-737, Physik Instrumente). The synchronized imaging process was controlled by a self-built LabVIEW (National Instruments) program. Supplementary Fig. S13 shows the optical layout and photographs of the system.

#### Macro-view Bessel-sheet microscope

Our custom macro-view Bessel-sheet microscope for mouse brain imaging was based on a tunable dual-side plane illumination and a zoomable detection. With splitting the collimated beam into 2 paths by a 50/50 beam splitter (CCM1-BS013/M, Thorlabs), two synchronized scanning Bessel sheets were generated at both sides of the sample using the similar way described above. A pair of long working distance objectives (Mitutoyo, 10×/0.28 or 5×/0.14) combined with tube lenses (ITL200, Nikon) were used to generate final dual-side Bessel sheets with illumination range of 1 or 4 mm (∼1.3 or 2.7 μm thickness) adapted to the detection FOV of 12.6× or 3.2× (1 or 4 μm lateral resolution) from a zoomable upright microscope (MVX10, Olympus). During image acquisition, the Bessel sheets were synchronized with the rolling shutter of sCMOS camera (Hamamatsu ORCA-Flash4.0 V2) at 20 fps for recording 3.2× LR images, or alternatively at 5 fps for recording 12.6× HR images. A motorized *xyz* stage (Z825B, Thorlabs combined with SST59D3306, Huatian) moved the sample in three dimensions to enable image stitching for the entire brain. A LabVIEW program was developed to synchronize the parts to automatically implement the beam scanning, image acquisition and tile-stitching in the proper order. Supplementary Fig. S12 shows the optical layout and photographs of the system.

#### Gaussian light-sheet microscope

Besides the Bessel-sheets, a SPIM path was also built for high-speed imaging of live *C. elegans* acting in microfluidic chamber. The 3.3 mm collimated lasers were first sent to a pair of cylindrical lenses (Thorlabs, LJ1695RM-A and LJ1567RM-A) to expand the beam height to ∼4 mm, which is sufficient to cover the length the chamber (3 mm). Then through another focusing cylindrical lens (Thorlabs, LJ1703RM-A), a Gaussian light-sheet with FWHM thickness of ∼10 μm and confocal range of ∼900 μm matched to the width of the chamber (0.8 mm) was finally generate to illuminate the *C. elegans*. The sample together with the microfluidic chip were moved by a motorized stage along *z* direction (Z825B, Thorlabs) for enabling fast 3D imaging.

### 3.2 Dual-stage-processed network (DSP-Net)

#### Network architecture

Our DSP-Net consists of a resolver that enhances the details and reduces noises for the low-quality LR images, and an interpolator that de-pixelates the intermediate output of the resolver by further extracting and permuting features. We adopted the deep back-projection network (DBPN)[34] as the resolver (Supplementary Fig. S1), for its outstanding feature extracting ability brought by alternating up-sampling and down-sampling operations. We modified all the layers to allow the processing of 3D input, and removed the last deconvolution layer to keep the image size of output unchanged. For the interpolator (Supplementary Fig. S2), residual dense network (RDN)[33] was used with structures also modified for 3D computation. To up-sample the 3D images in RDN interpolator, we further introduced a sub-voxel convolution layer (Supplementary Fig. S2), which was derived from previously reported efficient ESPCNN[38]. The DSP-Net implemented in Python using Tensorflow and Tensorlayer[39].

#### Network training and inference procedure

The DSP-Net training was based on semi-synthetic dataset. For mouse brain, the HR images of mouse brain were first experimentally obtained by 12.6× Bessel-sheet setup while the synthetic LR images were artificially generated by degrading the HR images using a model which simulates the optical blurring of microscope and pixelization of camera (details see Supplementary Note S2). For single-cell, the LR images were simply the diffraction-limited measurements under 60× Bessel-sheet. The corresponding HR images were generated by 3D-SRRF computation based on 30 stacks of diffraction-limited measurements. For live *C. elegans*, the synthetic HR and LR 3D images of microbeads were generated to represent the point-like neuronal signals. All abovementioned experiment used a directly down-sampled version of the HR data as the MR data, which still contains aliased high-resolution details that might be completely lost in the LR data. At the training stage, LR data were imported into the resolver of the DSP-Net, and sequentially convolved with the filters of each convolutional layer. The pixel-wise mean-square-errors (MSE) between the resolver’s outputs and the MR data, termed as the resolving loss, works as cost function of the parameters in each filter. A gradient descent method was used to efficiently minimize the resolving loss by optimizing the distribution of these parameters. The converged outputs of the resolver continue to flow into the interpolator, where the image size increases via extending and re-arranging feature maps extracted by the filters in the convolutional layers, and the following sub-voxel convolutional layers (Supplementary Fig. S2). At the 2^nd^ stage, the dual sub-nets are mutual-feedback. The minimization of MSE between the converged outputs of interpolator and the HR ground truths could lead to the optimization of the parameters not only in the 2^nd^-stage interpolator, but also those in the 1^st^-stage resolver. Finally, the well-trained DSP-Net with optimized parameters in both sub-nets was used to deduce superior-resolution 3D image stacks from the LR 3D inputs.

At the inference stage, the entire LR image stack were automatically subdivided into multiple blocks for being processed block by block (typically 100 × 100 × 100 voxels), owing to the memory limit of the GPUs. The super-resolved patches were automatically stitched together into a complete SR image stack automatically. Using a single RTX 2080 Ti GPU for computation, the reconstruction voxel rate was ∼10 megavoxels per second. The data implementation details were elucidated in Supplementary Table S1 and Supplementary Note S1.

### 3.3 Mouse brain experiment

#### Imaging

Neuron-tagged transgenetic whole mouse brain (Thy1-GFP-M) and vessel-labelled whole mouse brain (Alexa-647) were imaged using our custom macro-view Bessel-sheet microscope (Supplementary Fig. S12). For obtaining adequate HR image stacks for network training, we first imaged several brain sub-regions, such as isocortex, striatum, hippocampus, and cerebellum, using 12.6× detection combined with 1.3 μm-thickness Bessel-sheet illumination. The acquired 3D image of each sub-region encompassed 2048 × 2048 × 500 voxels across a 1.1 × 1.1 × 0.5 mm^3^ volume size. The total imaging time is ∼30 minutes for 20 regions with 5-fps acquisition rate and 200-ms exposure time. 10 patches of HR image (2048 × 2048 × 500 for each) were pre-processed to generate the HR-MR-LR pairs for network training.

Then the whole-brain imaging was implemented using 3.2× detection plus 2.7 μm-thickness Bessel-sheet illumination. We sequentially imaged 6 blocks (2 × 3) of the brain with each one containing 2048 × 2048 × 1250 voxels across a 4 × 4 × 5 mm^3^ volume size. With such a low-magnification setup, the total imaging time for a whole brain is down to ∼6 minutes with 20 fps acquisition rate. The 6 blocks were then stitched into an LR whole brain using Grid/collection stitching plugin of ImageJ. The stitched LR whole brain was taken as input for in-parallel restoration by the trained DSP-Net. Finally, a super-resolved whole brain output encompassing 3.2 teravoxels across 10 × 8 × 5 mm^3^ volume size was obtained. <u>The 12.6× HR images to be compared at the validation stage were also acquired at 20 fps with an exposure time of 50 ms.</u>

#### Speed and throughput calculation

The speed and throughput for imaging whole mouse brain is calculated by:

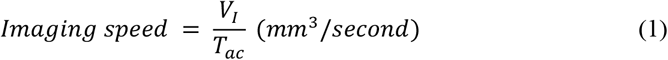

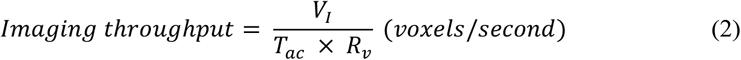

Where *V*_*I*_ is the imaging volume (mm^3^), *T*_*ac*_ is the acquisition time (second) and *R*_*v*_ is the volumetric resolution (μm^3^/voxel). We compared the achieved optical throughput for imaging a whole mouse brain by 3.2×, 6.4×, 12.6×, and DSP-Net-enabled 3.2× Bessel-sheets. To maintain an SNR high enough for distinguishing fine neuronal details, such as apical dendrites, the acquisition rate was 20 fps for 3.2× imaging, 10 fps for 6.4×, and 5 fps for 12.6× imaging. Under 3.2× setup, 6 tiles (2 × 3) with each one containing 2048 × 2048 × 1250 voxels across a 4 × 4 × 5 mm^3^ volume size were stitched together to form a whole brain dataset which finally encompassed 3891 × 5734 × 1250 voxels across 10 × 8 × 5 mm^3^ volume size. The total imaging time was around 6 minutes. For 6.4× and 12.6×, we stitched 20 (4 × 5) and 96 (8 × 12) tiles, respectively, with each one containing 7577 × 9420 × 2500 and 14950 × 22323 × 5000 voxels, respectively. The total imaging time was around 167 minutes for 6.4× acquisition and 1600 minutes for 12.6× acquisition. After the 4× resolution enhancement applied to 3.2× Bessel sheet, the achieved volumetric resolution was around 64 times higher than that from regular 3.2× Bessel-sheet, finally yielding ∼5 gigavoxels/s throughput which is over 200-times higher than that by 12.6× Bessel-sheet.

#### Neuron tracing

We chose a 0.5 × 0.4 × 0.2 mm^3^ volume in isocortex region to demonstrate the resolution enhancement from our DSP-Net can substantially benefit the neuron identification/tracing with improved accuracy. The data was converted and opened in Imaris software (Bitplane, USA). The Autopath mode of the Filament module was used to trace the same single neuron in LR, SR and HR images semi-automatically. We first assigned one point on the neuron to initiate the tracing. Then Imaris automatically calculated the pathway in accordance with the image data, reconstructed the 3D morphology and linked it with the previous part. This automatic procedure would repeat several times with manual adjustment involved until the neuron was finally correctly traced. Because the performance of the tracing highly relies on the resolution and SNR of the source image, the DSP-Net result allows obviously more accurate neuron tracing with finer fibers identified, as shown in Fig. 2d.

### 3.4 U2OS cell experiment

#### Imaging

The antibody-labeled microtubules and antibody-labeled endoplasmic reticulum in fixed *U2OS* cells were imaged by our custom high-magnification Bessel-sheet microscope. A thin Bessel-sheet provided high-axial-resolution optical sectioning (∼700 nm) of the cell, while a 60×/1.1 water dipping objective collected the fluorescence signals with a lateral resolution of ∼270 nm (pixel size 108 nm). During image acquisition, we repetitively scanned the cell using a piezo actuator (step size 300 nm) with excitation intensity ∼2 mW at the pupil plane of illuminating objective, obtaining 30 consecutive image stacks (2048 × 512 × 70 voxels for each stack) to reconstruct a super-resolution 3D image with a 3D SRRF computation procedure[32]. Then we used the SRRF results, the 2-time down-sampled SRRF results, and the diffraction-limited images of the 30^th^ acquisition as the HR, MR and LR data, respectively, for network training. 686 groups of such HR-MR-LR patch pairs with size 96 × 96 × 32 for each were included. For network validation, the same 60× Bessel sheet was applied to another *U2OS* cell with imposing regular-intensity excitation (∼2 mW), and low-intensity excitation (∼0.3 mW), to obtain LR, high-SNR and LR, low-SNR cell images, respectively.

#### Quantifications of spatiotemporal resolution and photon consumption

The fixed cell were imaged with a step size of 0.3 μm. Typically, when imaging adherent cells by a light-sheet fluorescence microscope, over 100 planes need to be recorded to form a 3D stack encompassing the entire cell. Limited by the response time of piezo stage, the volume acquisition rate was up to 2 Hz in this case, resulting in a total acquisition time of over 15 seconds for obtaining at least 30 volumes for 3D-SRRF of the cell. At the same time, the repetitive imaging also inevitably imposed increased photon burden to the sample. Using our DSP-Net, over 2-fold higher spatial resolution was achieved in all three dimensions, as compared to the diffraction-limited Bessel sheet imaging, with merely 1 volume of the cell quickly acquired in less than 1 seconds. From another point of view, it could surpass the diffraction limit with realizing 30-times higher temporal resolution, as compared to time series 3D-SRRF.

To ensure the reconstruction quality of 3D-SRRF, relatively high-intensity excitation was always required during the acquisition of the 30 image stacks. By contrast, DSP-Net permits significantly lower-dose excitation to the cell, owing to its strong capability for recovering very low-SNR image, which is obtained with an ∼7 fold lower-intensity excitation. Combined with the super resolution capability based on single measurement, the total photon consumption from DSP-Net can be over 2-orders-of-magnitude lower than that from the time series 3D-SRRF.

### 3.5 C. elegans experiment

#### Imaging

*C. elegans* were cultured on standard Nematode Growth Medium (NGM) plates seeded with OP50 and maintained at 22 °C incubators. The strain ZM9128 hpIs595[Pacr-2(s)::GCaMP6(f)::wCherry] of *C. elegans*, which expressed the calcium indicators to A-and B-class motor neurons, was used to detect the *Ca*^2+^ signaling in moving animals. For imaging neural activities, late L4 stage worms were loaded into a microfluidic chip with size of 3300 × 830 × 55 µm, allowing the worm to freely move within the FOV of a 4× objective (Olympus, XLFluor 4×/340, 0.28 NA, WD = 29.5 mm). During image acquisition, a 10 µm-thick Gaussian light-sheet illuminated the worm in the chamber from the side of the chip. The chip was mounted to a z-piezo stage, for being scanned at high speed. The sCMOS camera captured the plane images at a high speed of 400 fps (2048 × 512 pixel) and with a step size of 4.5 µm. The system as such permitted large-scale volumetric imaging (2048 × 512 × 13 voxel) at 30 volumes per second (VPS) acquisition rate across 3300 × 830 × 55 µm FOV. Due to the low phototoxicity provided by light-sheet illumination, we could continuously image the moving *C. elegans* for 30 seconds, recording the various worm behaviors such as forward, irregular and backward crawling, in the obtained 900 consecutive image stacks. The sequential 3D image stacks were then sent to the trained DSP-Net for be processed at high processing rate of 10-mega voxels per second by using a single RTX 2080 Ti graphic card.

#### Quantitative Analysis of *Ca*^*2+*^ dynamics and behavior of acting *C. elegans*

Following the previously reported method[40] we applied *Ca*^*2+*^ signal tracking to the 4D DSP-Net output. After the individual neurons were segmented in each image stack and automatically tracked throughout the entire 4D dataset, a manual correction was further applied to improve the accuracy of tracing. To extract the fluctuation of *Ca*^*2+*^ signals ΔF/F0, we calculated ΔF/F0 = (F(t)-F0)/F0 with F(t) being the fluorescence intensity of certain neuron at one time point, and F0 being the mean value of F(t) over the entire period. To analyze the worm behaviors in response to the neural activities, we first extracted the dynamic worm profiles using a U-Net based image segmentation[41]. Then the center lines of the worm were further extracted to calculate the time-varying curvatures of the worm[42], which were finally used to classify the worm behaviors as forward, backward and irregular crawling.

### 3.6 Cross sample experiments

A vessel-labelled whole mouse brain (Alexa-647) was imaged with the macro-view Bessel light-sheet microscope using the same 3.2× illumination and detection configurations mentioned above. Then the obtained image stack of vessels was recovered by the DSP-Nets trained with homogeneous vessel data and heterogeneous neuron data (Thy1-GFP-M).

Following the same logic, we also imaged the antibody-labeled endoplasmic reticulum in fixed *U2OS* cells (the antibody labeled EGFP) using the same conditions for imaging the microtubules. The diffraction-limited image stack of endoplasmic reticulum was super-resolved by DSP-Nets trained with endoplasmic reticulum data and microtubule data (Thy1-GFP-M).

### 3.7 Cross mode experiments

Coronal slices (100-μm thick) of a transgenetic mouse brain (Thy1-GFP-M) were imaged by a confocal microscope (Nikon Ni-E) using a 16×/0.8 water-dipping objective (CFI LWD 16×/0.8W). The voxel resolution was 0.31 × 0.31 × 1 μm in each acquired 3D image stack. For conducting a DSP-Net training based on confocal data, we then re-sampled the confocal stacks with matching their voxel resolution to those of 3.2× (2 × 2 × 4 μm voxel) and 12.6× (0.5 × 0.5 × 1 μm voxel) Bessel-sheet image stacks, generating synthetic 3.2× and 12.6× confocal images. The re-sampled 12.6× confocal image stacks were used as HR images for DSP-Net training. The LR images were the blurred, noised and down-sampled version of 12.6× images, while the MR images were directly down-sampled from 12.6× images. At validation stage, the 3.2× synthetic confocal image stack (down-sampled, blurred and noise-added) was recovered by both DSP-Nets trained with confocal and Bessel-sheet data.

We imaged antibody-labeled microtubules in fixed *U2OS* cells with a Nikon N-SIM microscope using an apochromatic oil immersion objective (Nikon, CFI Apo TIRF 100×/1.49 oil). The raw wide-field images were acquired under conditions of five pattern phases spaced by 2π/5, three pattern orientations spaced 60 degree apart, and a step size of 200 nm. Then the 3D SIM images were reconstructed based on the acquired wide-field images using a commercial software (NIS-Elements), and the diffraction-limited wide-field images were obtained simply by an average of the fifteen measurements at each plane.

### 3.8 Image quality evaluation

The normalized root mean square error (NRMSE) was used to indicate the pixel-wise difference between the SR image and the HR ground truth. These two images were first normalized to about the same intensity range [0, 1] by a percentile normalization, which ensured the difference of background wouldn’t affect the quantification. Then the NRMSE was computed as

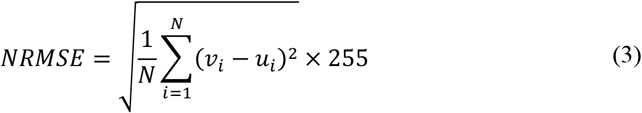

where N is the pixel number of the image (i.e. width × height), *v*_*i*_ and *u*_*i*_ are the *i*th pixel intensity of the SR and HR, respectively.

The signal-to-noise ratio (SNR) is computed as

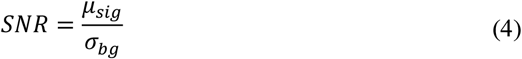

where *μ*_*sig*_ being the average intensity of the signal and *σ*_*bg*_ being the standard deviation of the background.

The resolution-scaled error (RSE) and resolution-scaled Pearson (RSP)[43] between the SR (or HR) images and the LR inputs were used to quantify the reliability of reconstruction. We obtained the averaged values based on the analysis of 30 randomly-selected regions using the Nanoj Squirrel ImageJ plugin.

### 3.9 Code availability

Code for DSP-Net and the examples are available at https://github.com/xinDW/DVSR.

## 4. DISCUSSION

In summary, the marriage of DSP-Net with cutting-edge 3D microscopy enables high spatiotemporal resolution volumetric imaging of various types of specimens far beyond the optical throughput limit. While LSFM is a powerful tool that can image samples in three dimensions at high speed and low phototoxicity, its throughput remains limited by the system optics, which are subject to limited NA, and it also affected by a certain degree of pixelization. In this study, the combination of our all-3D DSP-Net greatly pushes the limits of LSFM to achieve an ultra-high optical throughput that was previously unmet. Unlike current one-stage methods that directly map severely degraded LR measurements to HR targets, our DSP-Net divides the SISR problem into two sub-issues of de-blurring and de-pixelization that are solved successively, in effect fundamentally inverting the microscopy imaging processes that transform a high-resolution target into a blurred and pixelated image. This step-by-step strategy permits the network to extract more abundant features to create strong correlations between HR targets, MR aliased images, and degraded LR measurements, enabling a more accurate mapping of LR measurements to HR targets, which might be degraded in conventional one-stage deep learning networks. The experimental results verify that our DSP-Net can recover a 3D fluorescence image, with the DSP-Net super-resolving more details than other state-of-the-art super-resolution networks. The image resolution improvement achieved by the current DSP-Net is about four times in each dimension. It is possible to achieve a larger resolution-enhancement factor by combining progressive growing methods, provided that the equally accumulated artefacts can be suppressed at the same time. To implement this training strategy, a large number of 3D images captured at various magnifications are necessary to establish reliable targets for every growth stage of the progressive training.

The increased spatiotemporal information provided by the DSP-Net LSFM, together with its non-iterative rapid image reconstruction, are crucial for high-throughput imaging of large-size samples or the sustained recording of biological dynamics at high resolution. The advances made by the DSP-Net LSFM are demonstrated by the notably increased throughput for 3D high-resolution mapping of clarified large organs, the breaking of the diffraction limit for imaging of intracellular organelles with faster and lower phototoxicity measurements, and the improved temporal resolution for capturing instantaneous biological processes. Besides its combination with the LSFM, the DSP-Net has also shown its ability for appropriate cross-mode and cross-sample recovery, indicating extra flexibility allowing a network pre-trained on a relatively small amount of desired HR data from a relatively specialized system (e.g., fine neurons imaged by Bessel light-sheet) to be extensively applied to various types of samples showing similar structural characteristics (e.g., line-like vessels and microtubules) that were obtained using other imaging methods such as commercial confocal and wide-field microscopes, which may be more readily accessible. Therefore, we believe that DSP-Net is transformative, and that it could potentially bring new insights for wider application of the current computation-based imaging methods in biomedical research, which unceasingly requires ever-higher spatiotemporal performance.

## Acknowledgements

We thank Di Jin at MIT, Shaoqun Zeng and Xiang Bai at Huazhong University of Science &Technology for their helpful advices on the project design. We thank Yanyi Huang at Peking University and Jianbin Wang at Tsinghua University for their discussions and comments on the manuscript. We thank Yi Li, Yusha Li, Peng Wan, Tingting Yu, Dan Zhu, and Shangbang Gao for their help on the samples and biological analysis. Confocal experiment was done at the Wuhan National Laboratory for Optoelectronics at Huazhong University of Science and Technology and SIM experiment was done at the State Key Laboratory of Agricultural Microbiology, Huazhong Agricultural University. We thank Karl Embleton, PhD, from Liwen Bianji, Edanz Group China (www.liwenbianji.cn/ac), for editing the English text of a draft of this manuscript.

## Author contributions

P.F. and H.Z. conceived the idea. P.F., and H.Z. oversaw the project. C.F. and Y.Z. developed the optical setups. H.Z. developed the programs. C.F., and Y.Z. conducted the microfluidic and optical experiments. H.Z., C.F., Y.Z., and P.F. processed the images. H.Z., C.F., Y.Z., M.Z., Y.Z. and P.F. analyzed the data and wrote the paper.

## REFERENCES

1. J. B. Pawley, “Handbook of Biological Confocal Microscopy,” Journal of Biomedical Optics 25, 029902 (1995).

2. W. Denk, ., J. H. Strickler, and W. W. Webb, “Two-photon laser scanning fluorescence microscopy,” Science 248, 73–76 (1990).

3. H. Jan, S. Jim, D. B. Filippo, W. Joachim, and E. H. K. Stelzer, “Optical sectioning deep inside live embryos by selective plane illumination microscopy,” Science 305, 1007–1009 (2004).

4. R. H. Webb, “Confocal optical microscopy,” Reports on Progress in Physics 59, 427–471 (1996).

5. F. Helmchen and W. Denk, “Deep tissue two-photon microscopy,” Nature Methods 2, 932–940 (2005).

6. P. J. Keller, A. D. Schmidt, J. Wittbrodt, and E. H. Stelzer, “Reconstruction of zebrafish early embryonic development by scanned light sheet microscopy,” Science 322, 1065–1069 (2008).

7. M. B. Ahrens, M. B. Orger, D. N. Robson, J. M. Li, and P. J. Keller, “Whole-brain functional imaging at cellular resolution using light-sheet microscopy,” Nature Methods 10, 413 (2013).

8. P. J. Verveer, J. Swoger, F. Pampaloni, K. Greger, M. Marcello, and E. H. Stelzer, “High-resolution three-dimensional imaging of large specimens with light sheet-based microscopy,” Nature Methods 4, 311–313 (2007).

9. R. Tomer, K. Khairy, F. Amat, and P. J. Keller, “Quantitative high-speed imaging of entire developing embryos with simultaneous multiview light-sheet microscopy,” Nature Methods 9, 755 (2012).

10. J. G. Ritter, R. Veith, A. Veenendaal, J. P. Siebrasse, and U. Kubitscheck, “Light sheet microscopy for single molecule tracking in living tissue,” PLoS One 5, e11639–e11639 (2010).

11. P. J. Keller, A. D. Schmidt, A. Santella, K. Khairy, Z. Bao, J. Wittbrodt, and E. H. K. Stelzer, “Fast, high-contrast imaging of animal development with scanned light sheet–based structured-illumination microscopy,” Nature Methods 7, 637–642 (2010).

12. T. Chakraborty, M. K. Driscoll, E. Jeffery, M. M. Murphy, P. Roudot, B.-J. Chang, S. Vora, W. M. Wong, C. D. Nielson, H. Zhang, V. Zhemkov, C. Hiremath, E. D. De La Cruz, Y. Yi, I. Bezprozvanny, H. Zhao, R. Tomer, R. Heintzmann, J. P. Meeks, D. K. Marciano, S. J. Morrison, G. Danuser, K. M. Dean, and R. Fiolka, “Light-sheet microscopy of cleared tissues with isotropic, subcellular resolution,” Nature Methods 16, 1109–1113 (2019).

13. F. F. Voigt, D. Kirschenbaum, E. Platonova, S. Pagès, R. A. A. Campbell, R. Kastli, M. Schaettin, L. Egolf, A. van der Bourg, P. Bethge, K. Haenraets, N. Frézel, T. Topilko, P. Perin, D. Hillier, S. Hildebrand, A. Schueth, A. Roebroeck, B. Roska, E. T. Stoeckli, R. Pizzala, N. Renier, H. U. Zeilhofer, T. Karayannis, U. Ziegler, L. Batti, A. Holtmaat, C. Lüscher, A. Aguzzi, and F. Helmchen, “The mesoSPIM initiative: open-source light-sheet microscopes for imaging cleared tissue,” Nature Methods 16, 1105–1108 (2019).

14. G. Zheng, R. Horstmeyer, and C. Yang, “Wide-field, high-resolution Fourier ptychographic microscopy,” Nature Photonics 7, 739–745 (2013).

15. X. Ou, G. Zheng, and C. Yang, “Embedded pupil function recovery for Fourier ptychographic microscopy,” Optics Express 22, 4960–4972 (2014).

16. M. G. L. Gustafsson, “Surpassing the lateral resolution limit by a factor of two using structured illumination microscopy,” Journal of Microscopy 198, 82–87 (2010).

17. M. G. L. Gustafsson, “Nonlinear structured-illumination microscopy: wide-field fluorescence imaging with theoretically unlimited resolution,” Proceedings of the National Academy of Sciences of the United States of America 102, 13081–13086 (2005).

18. S. W. Hell and J. Wichmann,. “Breaking the diffraction resolution limit by stimulated emission: stimulated-emission-depletion fluorescence microscopy,” Optics Letters 19, 780–782 (1994).

19. M. J. Rust, M. Bates, and X. Zhuang, “Stochastic optical reconstruction microscopy (STORM) provides sub-diffraction-limit image resolution,” Nature Methods 3, 793 (2006).

20. B. Eric, G. H. Patterson, S. Rachid, L. O Wolf, O. Scott, J. S. Bonifacino, M. W. Davidson, L. S. Jennifer, and H. F. Hess, “Imaging intracellular fluorescent proteins at nanometer resolution,” Science 313, 1642–1645 (2006).

21. W. T. Freeman, T. R. Jones, and E. C. Pasztor, “Example-based super-resolution,” Computer Graphics & Applications IEEE 22, 56–65 (2002).

22. C. E. Duchon, “Lanczos Filtering in One and Two Dimensions,” J.appl.meteor 18, 1016–1022 (1979).

23. X. Li,. and M. T. Orchard, “New edge-directed interpolation,” IEEE Trans Image Process 10, 1521–1527 (2001).

24. K. Kwang In and K. Younghee, “Single-image super-resolution using sparse regression and natural image prior,” IEEE Transactions on Pattern Analysis & Machine Intelligence 32, 1127 (2010).

25. D. Weisheng, Z. Lei, S. Guangming, and W. Xiaolin, “Image deblurring and super-resolution by adaptive sparse domain selection and adaptive regularization,” IEEE Transactions on Image Processing 20, 1838–1857 (2011).

26. Y. Rivenson, Z. Göröcs, H. Günaydin, Y. Zhang, H. Wang, and A. Ozcan, “Deep learning microscopy,” Optica 4, 1437–1443 (2017).

27. H. Wang, Y. Rivenson, Y. Jin, Z. Wei, R. Gao, H. Günaydın, L. A. Bentolila, C. Kural, and A. Ozcan, “Deep learning enables cross-modality super-resolution in fluorescence microscopy,” Nature Methods 16, 103–110 (2019).

28. M. Weigert, U. Schmidt, T. Boothe, A. Müller, A. Dibrov, A. Jain, B. Wilhelm, D. Schmidt, C. Broaddus, S. Culley, M. Rocha-Martins, F. Segovia-Miranda, C. Norden, R. Henriques, M. Zerial, M. Solimena, J. Rink, P. Tomancak, L. Royer, F. Jug, and E. W. Myers, “Content-aware image restoration: pushing the limits of fluorescence microscopy,” Nature Methods 15, 1090–1097 (2018).

29. H. Zhang, C. Fang, X. Xie, Y. Yang, W. Mei, D. Jin, and P. Fei, “High-throughput, high-resolution deep learning microscopy based on registration-free generative adversarial network,” Biomedical Optics Express 10, 1044–1063 (2019).

30. E. Nehme, D. Freedman, R. Gordon, B. Ferdman, L. E. Weiss, O. Alalouf, T. Naor, R. Orange, T. Michaeli, and Y. Shechtman, “DeepSTORM3D: dense 3D localization microscopy and PSF design by deep learning,” Nature Methods 17, 734–740 (2020).

31. Y. Wu, Y. Rivenson, H. Wang, Y. Luo, E. Ben-David, L. A. Bentolila, C. Pritz, and A. Ozcan, “Three-dimensional virtual refocusing of fluorescence microscopy images using deep learning,” Nature Methods 16, 1323–1331 (2019).

32. R. Chen, Y. Zhao, M. Li, Y. Wang, L. Zhang, and P. Fei, “Efficient super-resolution volumetric imaging by radial fluctuation Bayesian analysis light-sheet microscopy,” Journal of Biophotonics 13, e201960242 (2020).

33. Y. Zhang, Y. Tian, Y. Kong, B. Zhong, and Y. Fu, “Residual Dense Network for Image Restoration,” in arXiv e-prints, (2018).

34. M. Haris, G. Shakhnarovich, and N. Ukita, “Deep Back-Projection Networks For Super-Resolution,” in arXiv e-prints, (2018).

35. Z. Wang, H. Zhang, L. Zhu, G. Li, Y. Li, Y. Yang, M. Roustaei, S. Gao, T. K. Hsiai, and P. Fei, “Network-based instantaneous recording and video-rate reconstruction of 4D biological dynamics,” bioRxiv, 432807 (2019).

36. T. Kawano, Michelle D. Po, S. Gao, G. Leung, William S. Ryu, and M. Zhen, “An Imbalancing Act: Gap Junctions Reduce the Backward Motor Circuit Activity to Bias *C.elegans* for Forward Locomotion,” Neuron 72, 572–586 (2011).

37. Q. Wen, S. Gao, and M. Zhen, “*Caenorhabditis elegans* excitatory ventral cord motor neurons derive rhythm for body undulation,” Philosophical Transactions of the Royal Society B: Biological Sciences 373, 20170370 (2018).

38. W. Shi, J. Caballero, F. Huszár, J. Totz, A. P. Aitken, R. Bishop, D. Rueckert, and Z. Wang, “Real-Time Single Image and Video Super-Resolution Using an Efficient Sub-Pixel Convolutional Neural Network,” in 2016 IEEE Conference on Computer Vision and Pattern Recognition (CVPR), 2016), 1874–1883.

39. H. Dong, A. Supratak, L. Mai, F. Liu, A. Oehmichen, S. Yu, and Y. Guo, “TensorLayer: A Versatile Library for Efficient Deep Learning Development,” in arXiv e-prints, (2017).

40. J. Y. Tinevez, N. Perry, J. Schindelin, G. M. Hoopes, G. D. Reynolds, E. Laplantine, S. Y. Bednarek, S. L. Shorte, and K. W. Eliceiri, “TrackMate: An open and extensible platform for single-particle tracking,” Methods, S1046202316303346.

41. T. Falk, D. Mai, R. Bensch, Ö. Çiçek, A. Abdulkadir, Y. Marrakchi, A. Böhm, J. Deubner, Z. Jäckel, K. Seiwald, A. Dovzhenko, O. Tietz, C. Dal Bosco, S. Walsh, D. Saltukoglu, T. L. Tay, M. Prinz, K. Palme, M. Simons, I. Diester, T. Brox, and O. Ronneberger, “U-Net: deep learning for cell counting, detection, and morphometry,” Nature Methods 16, 67–70 (2019).

42. C. Restif, C. Ibáñez-Ventoso, M. M. Vora, S. Guo, D. Metaxas, M. Driscoll, and A. Prlic, “CeleST: Computer Vision Software for Quantitative Analysis of C. elegans Swim Behavior Reveals Novel Features of Locomotion,” Plos Computational Biology 10, e1003702.

43. S. Culley, D. Albrecht, C. Jacobs, P. M. Pereira, C. Leterrier, J. Mercer, and R. Henriques, “Quantitative mapping and minimization of super-resolution optical imaging artifacts,” Nature Methods 15, 263 (2018).

